# Sequence-based deep learning model for annotating lytic polysaccharide monooxygenase families

**DOI:** 10.1101/2025.06.05.658061

**Authors:** Sweety Deena Ramesh, Kavin S Arulselvan, Vaishnavi Saravanan, Priyadharshini Pulavendran, Ragothaman M. Yennamalli

**Affiliations:** Department of Bioinformatics, School of Chemical and Biotechnology, SASTRA Deemed to be University, Thanjavur, Tamil Nadu, India 613401

**Author notes:** Corresponding Author: Dr. Ragothaman M. Yennamalli, 202, ASK2, Department of Bioinformatics, School of Chemical and Biotechnology, SASTRA Deemed to be University, Thanjavur, Tamil Nadu, India 613401. Equal contribution (Kavin S Arulselvan, Vaishnavi Saravanan, Priyadharshini Pulavendran).

**Keywords:** LPMO, Machine learning, Neural Network, Bi-LSTM, CAZymes

## Abstract

Lytic polysaccharide monooxygenases (LPMOs) are crucial enzymes that enhance the breakdown of polysaccharides, especially for biofuel production. Current computational tools for LPMO annotation are limited to a few families, leaving many unexplored. Here, we introduce PreDSLpmo v2.0, a deep learning-based tool that classifies and annotates the fast growing numbers of LPMOs across eight families using a curated dataset comprising over 30,000 LPMO sequences. To capture the compositional, physicochemical, and structural properties of these sequences, we extracted features using Python (iFeature) as well as R-based pipelines, generating over 13500 descriptors per sequence. Ensemble feature selection was used to identify significant features for binary and multiclass classification. To address data imbalance, we maintained a 1:1 ratio for all positive and negative sets during training and validation. A range of machine learning models were systematically trained and evaluated. An independent dataset was used to estimate the performance of the trained models. The multiclass Bi-LSTM model demonstrated the highest accuracy, robustness, and generalizability, outperforming feature-based approaches. We compared the performance of the model with Pfam, dbCAN3, and BlastP searches against the UniProtKB/swissprot database. The F1-score shows that the model’s predicted LPMO sequences are accurate. The reliability of predictions by Bi-LSTM model is comparable to that of dbCAN3 and often better, as confirmed by Pfam domain annotation and SignalP. The model, now deployed as a web server (https://predlpmo.in) for high-throughput, sequence-based functional annotation of LPMOs, provides a scalable and reliable solution for enzyme discovery in bioenergy and industrial biotechnology.

## Introduction

Lytic polysaccharide monooxygenases (LPMOs) are copper-dependent enzymes^1^ that oxidatively cleave glycosidic bonds in carbohydrate polymers, helping degrade abundantly available polysaccharides such as cellulose and chitin^2–3^. Cellulose and chitin, now discarded as waste has potential to be converted into biofuel and products useful in medicine and agriculture By catalyzing the cleavage of glycosidic bonds in crystalline polysaccharides, LPMOs enhance the accessibility of these substrates to hydrolytic enzymes, boosting overall saccharification efficiency.^4^ The ability to boost enzymatic hydrolysis has made LPMOs a focus of industrial and biotechnological research, particularly for the production of biofuels, sustainable energy sources that can help meet the rising energy demands while reducing dependence on fossil fuels.

Biofuel production involves the breakdown of plant biomass, especially crystalline polysaccharides such as cellulose, into fermentable sugars using enzymes. However, the stability of these polysaccharides under conventional physicochemical and enzymatic treatments presents a significant bottleneck for their industrial use, making the development of more effective biocatalysts a priority.^5^ Because LPMOs enable the oxidative break down of recalcitrant polysaccharide surfaces, they function as powerful biocatalysts for biomass degradation.

However, despite their importance for the biofuel industry, identifying and annotating LPMOs remains challenging, limiting the discovery and functional characterization of new enzymes. The systematic annotation of LPMO families is complicated due to their extensive sequence diversity.^6^ Despite this diversity, active-site configuration, including the characteristic histidine brace, is highly conserved across LPMOs.^7^ Conventional methods for identification and annotation are time-intensive and laborious.^8,9^ The semi-automated method to identify LPMOs using a sequence motif based on experimentally validated published literature, as implemented in the Carbohydrate-Active enZYme database, CAZy^10^, has limited ability to annotate family-level LPMO sequences.^11^ Initially classified as CBM33 in bacteria and GH61 in fungi,^12^ LPMOs are now grouped under the auxiliary activity (AA) class^13^ in the CAZy database and sub-classified into several families based on sequence and substrate specificity. The sequence diversity of LPMOs has expanded rapidly, with new members identified across bacteria, fungi, viruses, and eukaryotes, underscoring the need for accurate and scalable annotation tools.^4^

While experimental methods for LPMO identification are valuable, they are labor-intensive and not scalable enough to cope with the influx of sequence data generated by modern genomics. Computational resources, such as dbCAN3, have enabled automated CAZyme annotation using HMM profiles and other algorithms.^14,15^ However, their specificity for LPMO family classification remains limited as new families and sequence variants are discovered.^16–18^

To address these challenges, PreDSLpmo was developed. The tool employs traditional neural networks based on extracted sequence features as well as Bi-directional Long Short-Term Memory (Bi-LSTM) models that learn directly from sequence data.^19^ These approaches demonstrated superior predictive power over existing tools such as dbCAN2, particularly for classifying sequences into major LPMO families such as AA9 and AA10. However, PreDSLpmo was limited in scope to the AA9 and AA10 families. Moreover, the tool required command-line execution for prediction.

In this study, we present PreDSLpmo v2.0. The tool expands the scope of classification to the existing eight LPMO families. We built, trained and tested feature-based models such as stochastic gradient descent (SGD) and neural networks as well as the sequence based model, Bi-LSTM, based on protein data available on CAZy, JGI and GenBank. The best model, PreDSLpmo v2.0, expands the scope of earlier work. PreDSLpmo v2.0 could distinguish LPMOs from non-LPMO enzymes and identify the different classes to which they belong. PreDSLpmo v2.0 is now deployed as a user-friendly web platform, enabling high-throughput and accessible LPMO annotation for the research community. The tool has been rigorously trained on curated datasets with redundancy reduction and quality filtering. We have addressed the limitations of the previous approach and provide a scalable, accurate, and accessible solution for the functional annotation of LPMOs in the context of biofuel research and industrial biotechnology. We also introduce a web-server where PreDSLpmo v2.0 is now deployed as a user-friendly web platform, enabling high-throughput and accessible LPMO annotation for the research community.

## Methods

To screen and classify LPMO families, we developed PreDSLpmo v2.0, a machine learning-based tool. The tool uses multiclass prediction for the functional annotation of protein sequences. The sequences are classified as one of the existing eight LPMO families (AA9-AA11 and AA13-AA17) or as non-LPMO (AA1-AA8 and AA12).

The workflow, shown in Figure 1, began with the extraction of protein sequence data from the CAZy, NCBI and JGI (Joint Genome Institute) databases. The data extracted was preprocessed to ensure quality. Comprehensive feature extraction is then performed using iFeature and custom R scripts, generating 13,526 descriptors per sequence. These descriptors capture sequence composition and autocorrelation as well as physicochemical, and evolutionary features.

**Figure 1:**
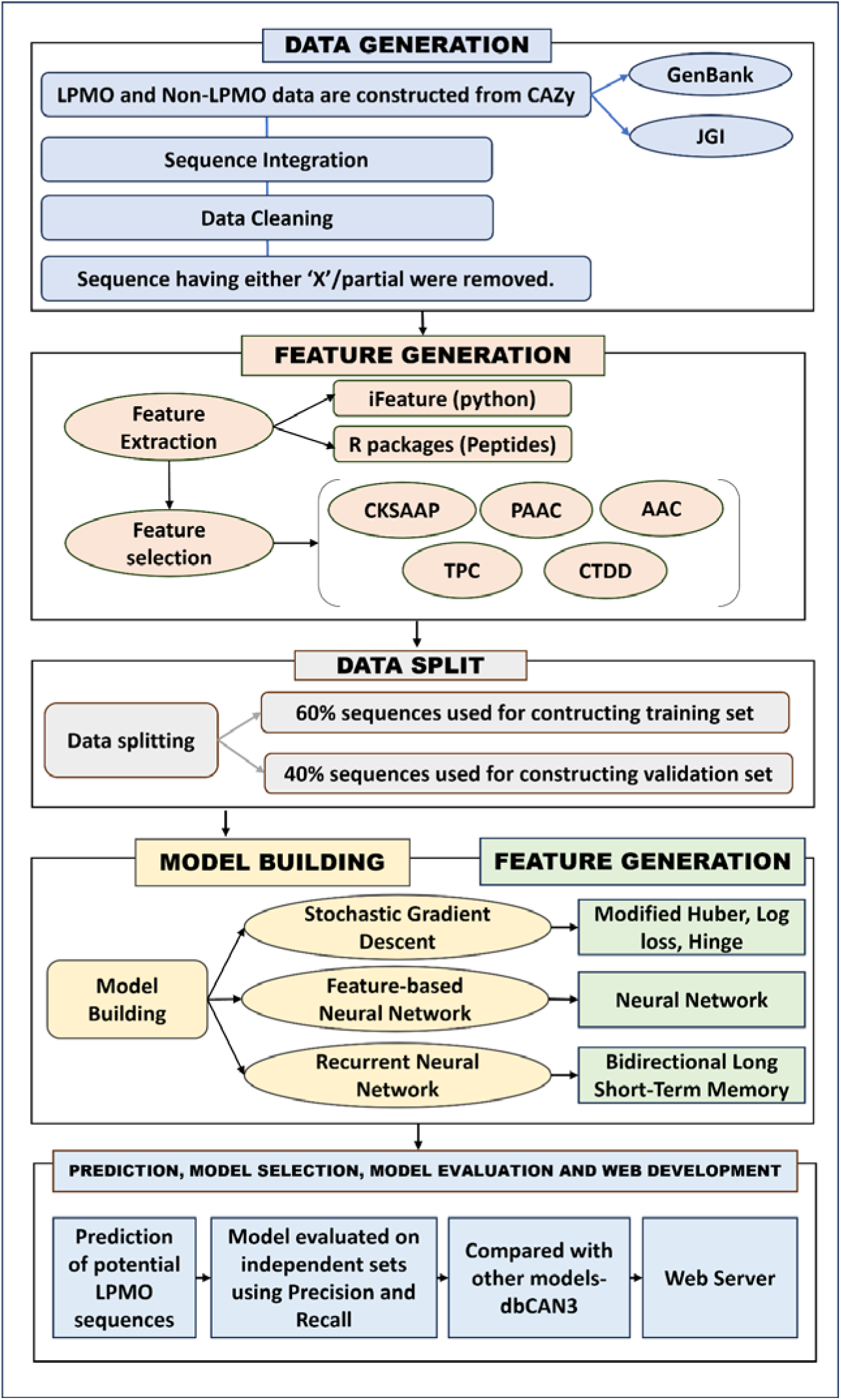
Workflow outlining the pipeline for automated LPMO family classification. Sequence data from GenBank and JGI were integrated, and cleaned to remove ambiguous or partial entries. Feature extraction was performed using Python (iFeature) as well as R packages, followed by feature selection and a 60:40 train-validation split. Machine learning models, SGD, neural networks, and Bi-LSTM, were trained and evaluated. The best-performing models were deployed as a user-friendly web server for high-throughput LPMO annotation.

The sequences of the proteins in the LPMO families, AA9-AA11 and AA13-AA17, as well as those in the non-LPMO families, AA1-AA8 and AA12, were downloaded from CAZy (accessed on 6th January, 2025). The corresponding EC numbers for each AA family are listed in Supplementary Table 1.

The sequences were retrieved from GenBank and JGI. From GenBank, sequences were extracted using accession IDs through the NCBI Entrez API, with a batch retrieval limit of up to 700 sequences per request.^20^ These sequences were downloaded in FASTA format. The JGI portal was accessed to download sequences based on organism names and gene IDs. Initial retrieval involved queries using the names of organisms, followed by fetching specific genome sequences corresponding to the identified organisms. A Python script was employed to automate sequence retrieval after downloading organism data. The CAZy database in JGI contained repeated entries with the same sequence ID and organism, leading to redundancy. A Python script was used to remove duplicates. Sequences containing ambiguous residues (’X’) were eliminated, and sequences labeled as partial in the header were excluded using the Python script to maintain dataset integrity.

For clustering and removing redundant protein or nucleotide sequences, Cluster Database at High Identity with Tolerance (CD-HIT) is a widely used bioinformatics tool.^21^ We initially used CD-HIT with a 65% sequence identity threshold for the positive dataset (AA9 to AA17, except the AA12 families of LPMOs) and observed that 52% to 91% of the sequences were culled. When the cutoff was kept at 70%, only 48% to 86% of the sequences were culled. Similarly, for the negative dataset, we observed that 73% to 86% of the sequences were culled at the cutoff of 65 % sequence identity. For the 70% cutoff, we observed that 67% to 81% of sequences were culled. To maintain a balanced dataset with a diverse set of sequences, we settled on a 70% sequence identity cutoff to remove redundant sequences. This cutoff ensured the formation of balanced training and validation sets across the AA1 to AA17 families. The cleaned dataset as well as the CD-HIT-filtered dataset were used to train the model.

**Table 1:** Sequence counts for LPMO family for training and validation set. The number of sequences downloaded from each database, cleaned, and combined for each AA family.

| LPMO Family | Count as per CAZy Database* | Redundancy-removed sequences | Genbank Downloaded | JGI Downloaded | Genbank Cleaned | JGI Cleaned | Genbank and JGI Combined | CD Hit** |
| --- | --- | --- | --- | --- | --- | --- | --- | --- |
| AA9 | 15106 | 15088 | 1393 | 13673 | 1321 | 13649 | 14970 | 3853 |
| AA10 | 12337 | 12332 | 12281 | 51 | 11528 | 51 | 11579 | 1336 |
| AA11 | 3482 | 3480 | 381 | 3074 | 370 | 3072 | 3442 | 1541 |
| AA13 | 414 | 414 | 75 | 338 | 65 | 338 | 403 | 56 |
| AA14 | 1393 | 1393 | 143 | 1249 | 142 | 1246 | 1388 | 513 |
| AA15 | 742 | 735 | 552 | 183 | 490 | 183 | 673 | 346 |
| AA16 | 1094 | 1093 | 106 | 987 | 104 | 986 | 1090 | 328 |
| AA17 | 161 | 157 | 88 | 69 | 77 | 69 | 146 | 75 |

\*-Accessed January 2025
\*\* -70% Identity

**Table 2:**
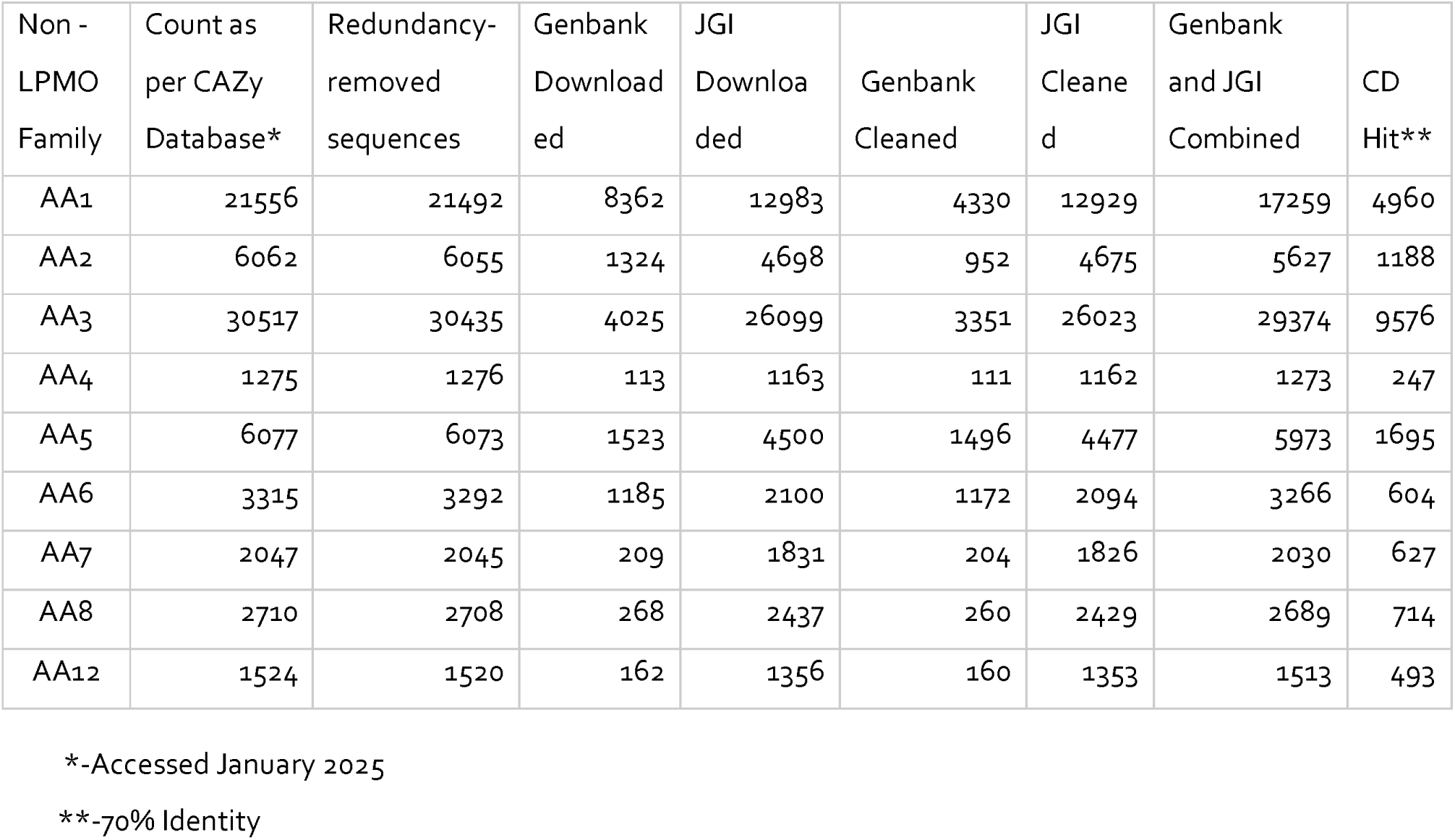
Sequence counts for non-LPMO family for training and validation set. The number of sequences downloaded from the GenBank and JGI databases, cleaned, and combined for each AA family.

| Non - LPMO Family | Count as per CAZy Database* | Redundancy-removed sequences | Genbank Downloaded | JGI Downloaded | Genbank Cleaned | JGI Cleaned | Genbank and JGI Combined | CD Hit** |
| --- | --- | --- | --- | --- | --- | --- | --- | --- |
| AA1 | 21556 | 21492 | 8362 | 12983 | 4330 | 12929 | 17259 | 4960 |
| AA2 | 6062 | 6055 | 1324 | 4698 | 952 | 4675 | 5627 | 1188 |
| AA3 | 30517 | 30435 | 4025 | 26099 | 3351 | 26023 | 29374 | 9576 |
| AA4 | 1275 | 1276 | 113 | 1163 | 111 | 1162 | 1273 | 247 |
| AA5 | 6077 | 6073 | 1523 | 4500 | 1496 | 4477 | 5973 | 1695 |
| AA6 | 3315 | 3292 | 1185 | 2100 | 1172 | 2094 | 3266 | 604 |
| AA7 | 2047 | 2045 | 209 | 1831 | 204 | 1826 | 2030 | 627 |
| AA8 | 2710 | 2708 | 268 | 2437 | 260 | 2429 | 2689 | 714 |
| AA12 | 1524 | 1520 | 162 | 1356 | 160 | 1353 | 1513 | 493 |

\*-Accessed January 2025
\*\*-70% Identity

To test the accuracy of the models for predicting the class of LPMO even when confronted with independent sets for AA9-AA11 and AA13-AA17, we downloaded new sequences reported to be AA1-AA17 by CAZy as of May 8th, 2025. The counts of new AA1-AA17 sequences which were added as independent sets were cleaned (Table 3 and Table 4). The increase in the number of LPMO sequences over a short time period highlights the growing interest of the research community in LPMO discovery and annotation.

**Table 3:**
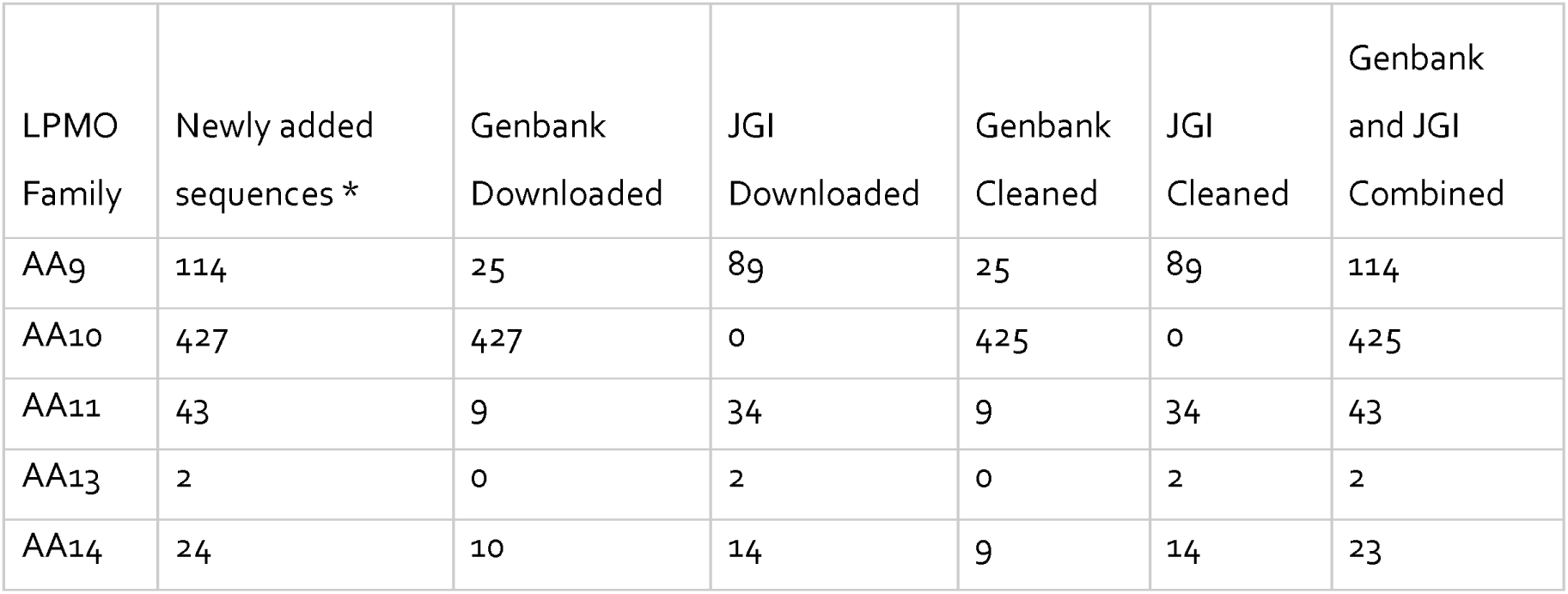
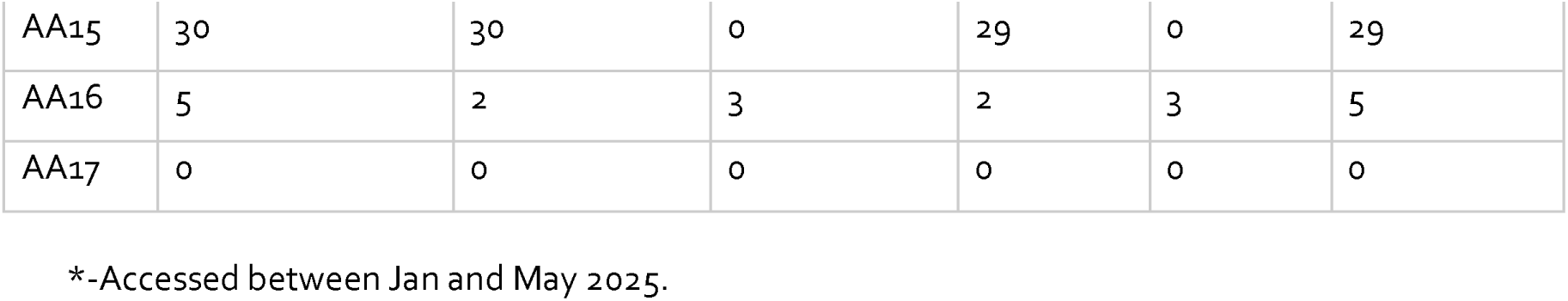
Sequence counts for LPMO family for independent sets. The number of sequences downloaded from the GenBank and JGI databases, cleaned, and combined for each AA family.

| LPMO Family | Newly added sequences * | Genbank Downloaded | JGI Downloaded | Genbank Cleaned | JGI Cleaned | Genbank and JGI Combined |
| --- | --- | --- | --- | --- | --- | --- |
| AA9 | 114 | 25 | 89 | 25 | 89 | 114 |
| AA10 | 427 | 427 | 0 | 425 | 0 | 425 |
| AA11 | 43 | 9 | 34 | 9 | 34 | 43 |
| AA13 | 2 | 0 | 2 | 0 | 2 | 2 |
| AA14 | 24 | 10 | 14 | 9 | 14 | 23 |
| AA15 | 30 | 30 | 0 | 29 | 0 | 29 |
| AA16 | 5 | 2 | 3 | 2 | 3 | 5 |
| AA17 | 0 | 0 | 0 | 0 | 0 | 0 |

\*-Accessed between Jan and May 2025.

**Table 4:** Sequence counts for non-LPMO family for independent sets. The number of sequences downloaded from the GenBank and JGI databases, cleaned, and combined for each AA family.

| Non -<br>LPMO<br>Family | Newly added<br>sequences * | Genbank<br>Downloaded | JGI<br>Downloaded | Genbank<br>Cleaned | JGI<br>Cleaned | Genbank and<br>JGI<br>Combined |
| --- | --- | --- | --- | --- | --- | --- |
| AA1 | 345 | 290 | 55 | 273 | 55 | 328 |
| AA2 | 97 | 84 | 13 | 82 | 13 | 95 |
| AA3 | 264 | 165 | 99 | 113 | 99 | 212 |
| AA4 | 3 | 2 | 1 | 2 | 1 | 3 |
| AA5 | 125 | 106 | 19 | 105 | 19 | 124 |
| AA6 | 153 | 144 | 9 | 131 | 9 | 140 |
| AA7 | 17 | 7 | 10 | 7 | 10 | 17 |
| AA8 | 13 | 6 | 7 | 6 | 7 | 13 |
| AA12 | 11 | 4 | 7 | 4 | 7 | 11 |

\*-Accessed between Jan and May 2025.

### Feature extraction

For feature extraction, to accurately represent the properties of protein sequences, we used a combination of Python and R-based tools. In Python, we used the iFeature package, which allowed us to extract 21 different types of numerical descriptors, resulting in 13,494 features per sequence.^22^ These features, covering a wide range of sequence characteristics, are mentioned in Supplementary Table 2. This approach provided a detailed, high-dimensional representation of each protein sequence.

Apart from the iFeature package, custom scripts in R were developed using the peptides package to generate six additional features for each sequence.^23^ These features focused on important biochemical properties such as molecular weight, isoelectric point, hydrophobicity, aliphatic index, instability index, and boman index.

By combining the comprehensive feature extraction capabilities of Python with specialized biochemical profiling in R, we ensured that our dataset captured both the overall patterns and the specific physicochemical properties of the sequences. This robust representation was essential for accurately distinguishing LPMO from non-LPMO protein families.

### Feature selection

To optimize inputs for machine learning models, ensemble feature selection protocols was implemented using the R package, ensemble feature selection (EFS).^24^ However, due to significant computational limitations, including long runtimes and memory allocation errors, this approach proved impractical for large, high-dimensional datasets. To address this challenge, we re-engineered the feature selection pipeline in Python to replicate and integrate six commonly used methods from R, including the Mann-Whitney U test (median-based), Pearson and Spearman correlations, Logistic Regression, and Random Forest classifiers using both entropy and Gini impurity criteria. Each method independently computed feature importance scores, which were then normalized and combined into a single EFS score, enabling improved efficiency and scalability.

For binary classification, the EFS framework that was applied combined six methods: median-based Mann–Whitney U test, Pearson and Spearman correlation analyses, Logistic Regression coefficients, and Random Forest importance scores using both Gini and entropy criteria. The feature rankings from these methods were normalized and aggregated into unified EFS scores (ranging from 0 to 1), allowing us to retain only the most discriminative features for each binary task.

For multiclass classification across nine classes (eight LPMO families and the non-LPMO group), we adapted a modified EFS protocol using multiclass-compatible metrics: Kruskal–Wallis H-test, ANOVA F-statistics, Mutual Information, Multinomial Logistic Regression, and Random Forest importance measures. This approach ensured the selection of features with strong discriminatory power across the nine classes, minimizing family-specific biases.

By leveraging Python’s computational libraries (*scikit-learn*, *scipy*, *numpy*), we achieved substantial improvements in runtime and memory use, reducing processing time by over 80% compared to the original R-based implementation. This dual strategy ensured robust and scalable feature selection, providing high-quality inputs for both binary and multiclass classification models.

### Data split

To prepare data for model training and evaluation, we divided each dataset into training and validation sets using a 60:40 split. Specifically, 60% of the sequences from both positive (LPMO) and negative (non-LPMO) classes were used for training, while the remaining 40% was set aside for validation. In both sets, we maintained a 1:1 ratio of positive and negative samples to ensure balanced class representation in Table 5.

**Table 5:** Data splitting of curated dataset for multiclass model approach.

| Positive set (LPMO) |  |  | Negative set (Non LPMO) |  |  |
| --- | --- | --- | --- | --- | --- |
| Family | Training | Validation | Family | Training | Validation |
| AA9 | 8982 | 5988 | AA1 | 5057 | 3371 |
| AA10 | 6947 | 4632 | AA2 | 1648 | 1099 |
| AA11 | 2065 | 1377 | AA3 | 8606 | 5737 |
| AA13 | 242 | 161 | AA4 | 373 | 248 |
| AA14 | 833 | 555 | AA5 | 1750 | 1166 |
| AA15 | 404 | 269 | AA6 | 956 | 638 |
| AA16 | 654 | 436 | AA7 | 595 | 396 |
| AA17 | 88 | 58 | AA8 | 787 | 525 |
|  |  |  | AA12 | 443 | 296 |
| Total | 20215 | 13476 | Total | 20215 | 13476 |

For binary classification models, we applied ensemble feature selection (EFS) thresholds to filter features before splitting the data. We created separate datasets by selecting features with EFS scores greater than 0.5 and, more stringently, greater than 0.6. This was done for both the cleaned dataset and the CD-HIT clustered dataset.

For the multiclass classification model (covering eight LPMO families and the non-LPMO group), we selected features with EFS scores above 0.5 for both cleaned and CD-HIT datasets. After feature selection, these datasets were also split into training and validation sets using the 60:40 ratio, ensuring balanced representation of all nine classes. This approach allowed us to systematically evaluate the impact of different feature selection thresholds and dataset types on model performance, while maintaining rigorous and unbiased data partitioning.

### Model building

Training and validation datasets derived from both cleaned and CD-HIT filtered sequences were used as input files into the developed Python scripts. The scripts implemented various machine learning algorithms from scikit-learn,^25^ keras, tensorflow package and machine learning libraries from Python.

Models were developed for the binary as well as the multiclass classification of LPMO families. Two models were feature-based: the stochastic gradient descent (SGD) classifier and feature-based neural networks. The third model, Bi-LSTM, was used to directly learn patterns from raw protein sequences.

For the binary classification, classifiers were trained for each LPMO family (AA9-AA11 and AA13-AA17). Within the feature-based framework, the SGD classifier was trained using three different loss functions, log loss, hinge loss, and modified Huber loss, resulting in three SGD-trained classifiers per family. The feature-based neural network contributed one additional classifier per family. This resulted in 32 classifiers for each feature-based approach across eight LPMO families. Overall, the SGD approach produced 96 binary classifiers (32 per loss function), and the feature-based neural network produced 32 binary classifiers. In contrast, the sequence-based Bi-LSTM architecture generated 16 binary classifiers, reflecting its direct use of raw sequence inputs rather than feature subsets.

All models were trained on both cleaned and CD-HIT reduced datasets and were evaluated using ensemble feature selection (EFS) thresholds of 0.5 and 0.6. Comparative analysis demonstrated that an EFS threshold of 0.5 consistently delivered optimal performance, which was subsequently adopted for all multiclass classification experiments.

Consistent with the binary classification strategy, multiclass models were trained on both cleaned and CD-HIT reduced datasets, to classify sequences into eight LPMO families and a non-LPMO class.

### Stochastic gradient descent (SGD)-based classification

Stochastic gradient descent (SGD) is an effective optimization algorithm to train linear classifiers on high-dimensional biological sequence data. It iteratively updates model weights to minimize differentiable loss functions, making it especially suitable for large-scale learning tasks. The SGD framework was evaluated using log loss, hinge loss, and modified Huber loss, to determine their relative effectiveness in classifying the LPMOs.

Log loss corresponds to logistic regression and provides probabilistic predictions, making it appropriate for multiclass classification. Hinge loss, commonly used in support vector machines (SVMs), emphasizes maximum-margin classification and is primarily used in binary classification. Modified Huber loss offers a robust, smooth alternative to hinge loss by combining margin-based classification with squared loss behavior, resulting in improved tolerance to outliers while enabling probability estimation.

To implement this approach, two separate pipelines were developed. For the multiclass classification, a single SGD classifier was trained. This model was designed to categorize sequences into one of nine classes: eight representing the LPMO families (AA9 to AA17) and one consolidated non-LPMO class. The dataset was preprocessed through imputation and normalization using SimpleImputer and StandardScaler. These preprocessing components were serialized using Joblib for future reproducibility. Hyperparameter tuning was done using GridSearchCV, evaluating a range of values for max_iter (50 to 500) and alpha (1e^-7^ to 1.0) across five-fold cross-validation. Class imbalance was handled by applying class weights and ElasticNet regularization. Early stopping and adaptive learning rates were included to stabilize training and prevent overfitting.

The core training logic resided in the function getBestModel(), which selected the optimal model based on cross-validation results. A heatmap of cross-validation accuracy was generated to visualize performance across hyperparameter combinations. Model evaluation and result generation were handled by saveBestModel(), which produced and saved a comprehensive set of outputs, including confusion matrices, classification reports, ROC curves, and Precision-Recall (PR) curves.

For multiclass classification, the script additionally employed label binarization to calculate class-wise ROC and PR metrics. Notably, the final trained model was also evaluated on the independent test set (previously unseen data), ensuring an unbiased assessment of the model’s generalizability and robustness.

For the binary classification, a separate model was built for each LPMO family. The corresponding top-feature dataset for that family was then loaded for both training and validation. As with the multiclass approach, preprocessing steps were applied and saved, and hyperparameters were optimized using grid search. The same three loss functions were explored by adjusting the loss parameter in the SGD classifier. The output was evaluated using saveBestModel(), which saved binary confusion matrices, classification reports, and PR/ROC plots tailored to each family and loss function combination. As in multiclass classification, each binary classification model was evaluated on the independent test set to validate performance beyond the training data.

The SGD-based classification pipeline provided a highly modular, interpretable, and computationally efficient alternative to deep learning architectures. The ability to swap out loss functions and apply models across both binary and multiclass tasks made this approach particularly flexible for benchmarking LPMO classification. Additionally, the inclusion of independent test set evaluations strengthened the reliability of the model results and their potential for real-world application in functional annotation workflows.

### Feature-based neural network

The neural network architectures comprised multiple dense layers. To enable accurate learning from high-dimensional sequence-derived features, adaptive optimization was used.

The sequences of each LPMO family were distinguished from non-LPMO sequences using individual binary classifiers constructed for the eight LPMO families (Figure 2A). The classifiers were implemented using a feedforward neural network with three hidden layers followed by an output layer. The network architecture comprised two hidden layers with 30 neurons each and a third hidden layer with 15 neurons. All hidden layers had a Rectified Linear Unit (ReLU) activation function to introduce non-linearity, enhancing the model’s capacity to learn complex patterns. The output layer contained two neurons, representing the target LPMO family and the corresponding non-LPMO class, with a softmax activation function to yield class probabilities. The models were trained for 100 epochs with a batch size of 50 using the Adam optimizer and sparse categorical cross-entropy loss function. To evaluate performance, a series of plots and metrics were generated, including training/validation accuracy curves, confusion matrices, Receiver Operating Characteristic (ROC) and Precision-Recall (PR) curves, as well as a feature correlation heatmap. This approach enabled robust binary discrimination for each family, and the models were independently saved along with classification reports and graphical outputs for interpretability and future deployment.

**Figure 2:**
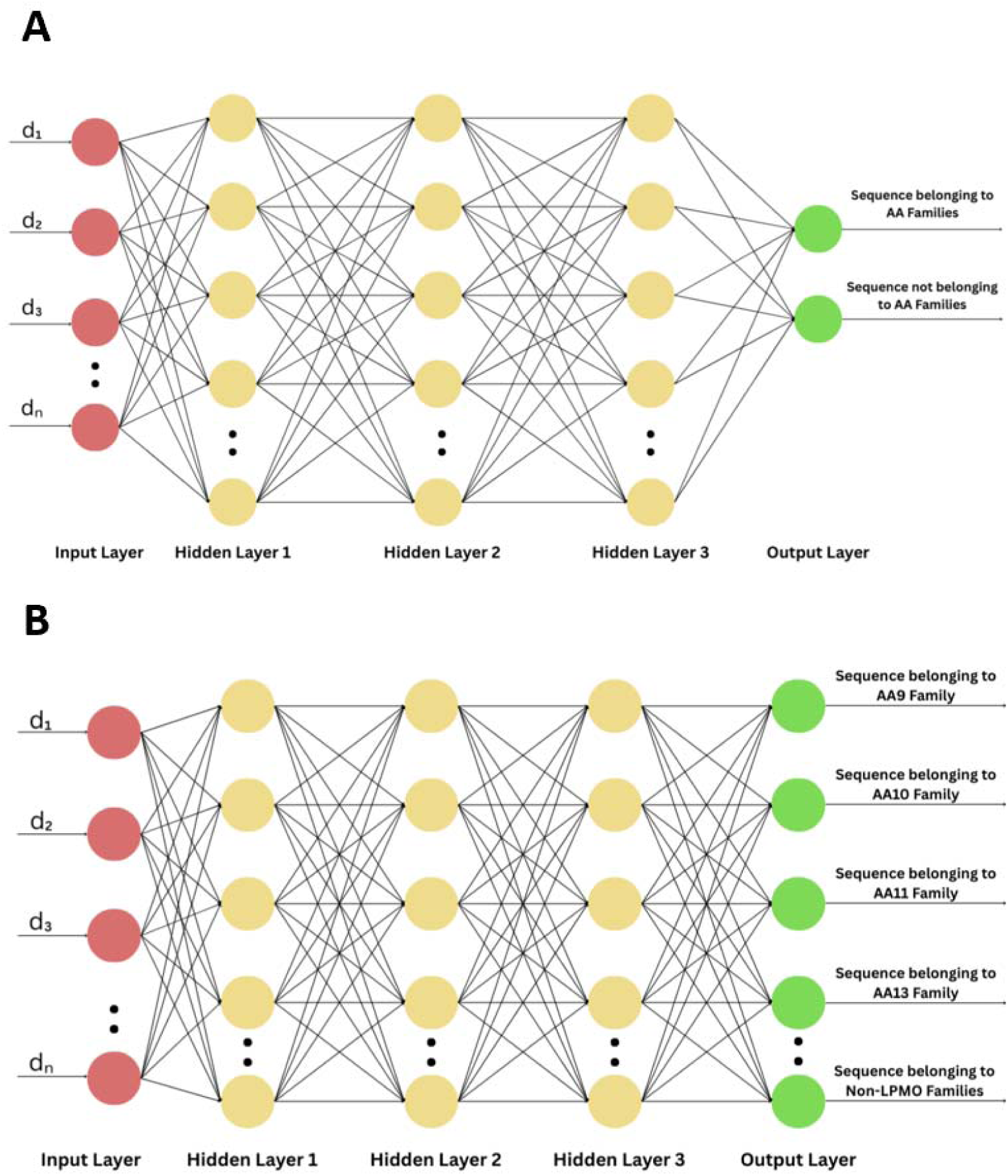
Schematic representation of feature-based neural network architectures used for LPMO family classification. (A) Binary neural network model distinguishing AA from non-AA sequences. (B) Multiclass neural network model classifying sequences into individual LPMO families (only four representative LPMO families are shown for simplicity) and non-LPMO class.

To develop a unified model capable of simultaneously identifying multiple LPMO families, we constructed a multiclass classification neural network (Figure 2B). The model was trained to classify input sequences into one of nine classes: eight corresponding to the LPMO families (AA9-AA17) and one consolidated non-LPMO class.

The model architecture consisted of four dense layers, including three hidden layers with ReLU activation and an output layer with softmax activation. The hidden layers contained 128, 64, and 32 neurons with dropout regularization (0.5 and 0.3). Batch normalization was applied to improve generalization. The output layer comprised nine neurons, each representing one class. To address class imbalance in the training dataset, class weights were computed and applied during training. The model was trained for a maximum of 200 epochs with a batch size of 128, incorporating early stopping and learning rate reduction on plateau as callbacks to prevent overfitting and dynamically adjust learning rates.

Comprehensive post-training analyses included confusion matrices, classification reports, and class-wise sample distribution assessments. The training and validation curves for accuracy and loss were also plotted to visualize learning dynamics.

### Bidirectional long short-term memory (Bi-LSTM)

We used Bi-LSTM networks to capture contextual relationships embedded within protein sequences. Bi-LSTM networks are proficient at processing sequential data. Bi-LSTM models leverage forward and backward sequence information, allowing them to learn patterns essential for effective classification. In this study, two configurations of Bi-LSTM networks were used: binary models trained to identify individual LPMO families, and a unified multiclass model developed to classify sequences across all eight LPMO families as well as a consolidated non-LPMO class.

For binary classification, a dedicated Bi-LSTM model was trained for each LPMO family (Figure 3A). Each model began with a character-level tokenization of protein sequences, followed by padding to a fixed sequence length to ensure input uniformity. The architecture consisted of an embedding layer with 300 dimensions, followed by a Bi-LSTM layer comprising 400 units. To enhance generalization and mitigate overfitting, dropout and recurrent dropout mechanisms were applied, both set at a rate of 0.5. The network had two fully connected layers with 100 and 50 neurons. ReLU activation functions were applied to both layers. To reduce overfitting, L2 regularization was applied to these layers. The final layer contained a single neuron activated by a sigmoid function to perform binary classification.

**Figure 3:**
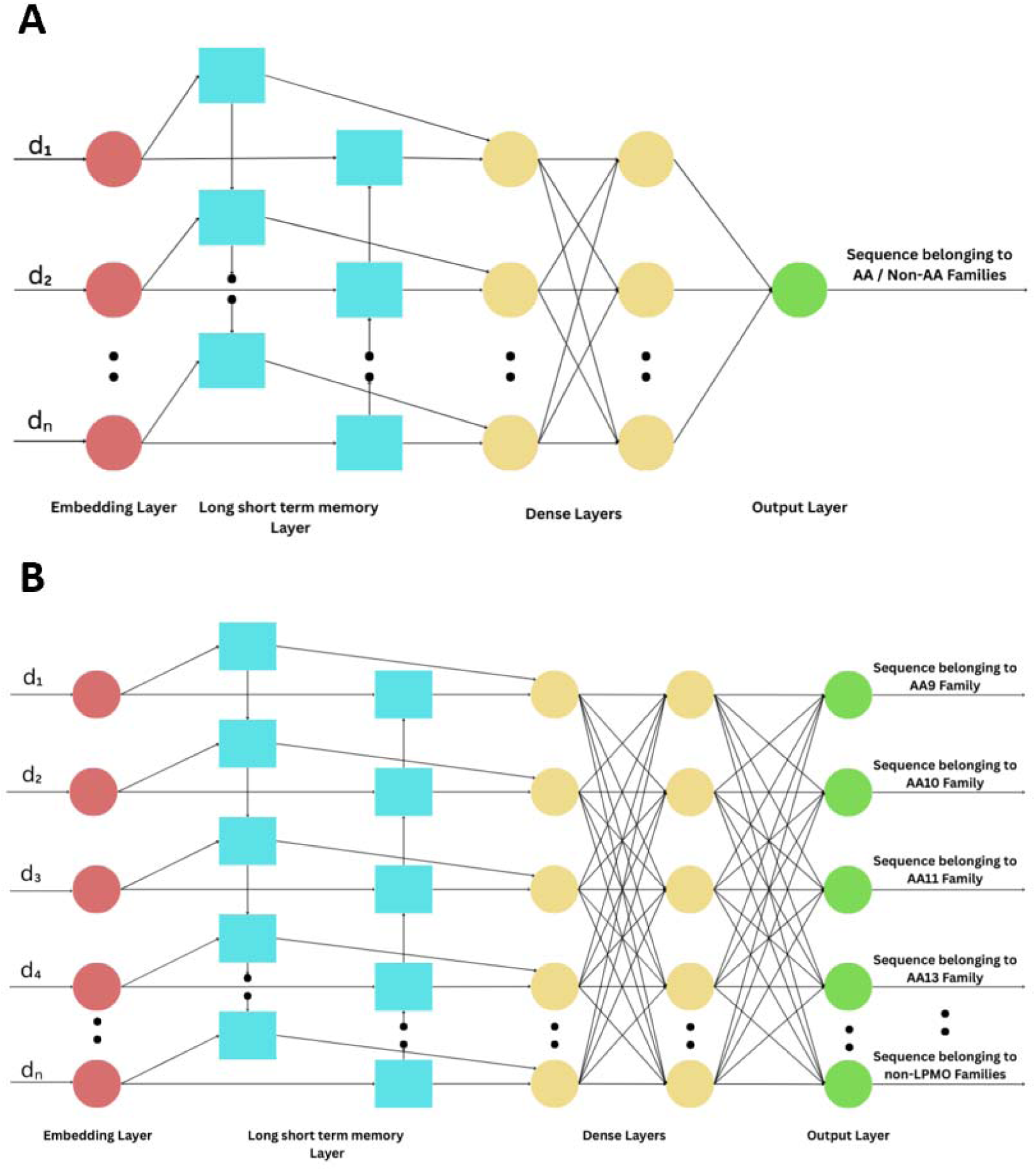
Schematic representation of Bi-LSTM architectures used for LPMO family classification. (A) Binary Bi-LSTM model classifying AA vs. non-AA sequences. (B) Multiclass Bi-LSTM model differentiating between LPMO families (Only four representative LPMO families are shown for simplicity) and non-LPMO class.

The models were trained using the Adam optimizer and binary cross-entropy loss. To account for class imbalance, class weights were dynamically computed based on the distribution of classes in the training data. Training was conducted for up to 100 epochs with a batch size of 128, employing early stopping and adaptive learning rate scheduling to ensure optimal convergence.

The performance of each binary model was evaluated using ROC and Precision-Recall curves, confusion matrices, and training history visualizations. This approach enabled high-sensitivity and family-specific predictions, making it well-suited for fine-grained LPMO detection.

The multiclass Bi-LSTM model was developed to classify sequences into one of nine classes: eight corresponding to individual LPMO families and one representing the non-LPMO group (Figure 3B). For this model, we used the same character-level sequence representation and padding approach as described above. The network architecture included a 256-dimensional embedding layer, followed by two stacked Bi-LSTM layers with 512 and 256 units, respectively. Regularization was applied using dropout layers and layer normalization to enhance stability and generalization during training. A dense layer with 256 neurons and ReLU activation was added before the final output layer. The output layer contained nine neurons activated by a softmax function to produce probability distributions across the target classes.

The model was trained using sparse categorical cross-entropy and the Adam optimizer, with class weights adjusted to handle imbalanced data. Training was conducted over a maximum of 200 epochs with a batch size of 256, using early stopping, learning rate reduction on plateau, and model checkpointing to retain the best-performing model state.

Evaluation of the multiclass model included per-class ROC curve analysis, confusion matrices, classification reports, and plots depicting the evolution of training accuracy and loss over epochs.

### Evaluation metrics

To rigorously evaluate classifier performance, we prioritized precision, recall, and F1-score as primary metrics, complemented by ROC-AUC and confusion matrices.^26^ These metrics were selected to address potential class imbalances and ensure robust interpretation of model efficacy.

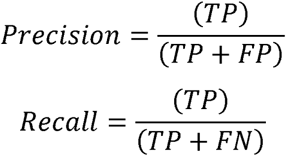

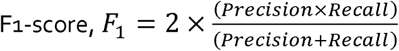

where, *TP* stands for True Positives, *FP* stands for False Positives, and *FN* stands for False Negatives.

For model validation, the performance of the trained Bi-LSTM model was assessed using an independent test set that was not seen during training. We first loaded the optimized Bi-LSTM along with its tokenizer. The tokenizer then converted sequences from the independent dataset into integer encoded formats, which were padded or truncated to a fixed length of 400 residues to match the model’s input requirements. The model generated predictions for each sequence, providing both class probabilities and the most likely class label across all nine categories (eight LPMO families and non-LPMO class).

To comprehensively evaluate the model’s predictive accuracy, a classification report was generated, detailing precision, recall, and F1-score for each class. Additionally, a confusion matrix was constructed and visualized as a heatmap to illustrate the distribution of true versus predicted labels. All results, including the classification report, confusion matrix, and the predicted class assignments, were saved for further analysis.

## Results and discussion

### Dataset curation

We obtained sequences from the NCBI and JGI databases based on the entries given in CAZy. The total number of sequences downloaded for each family (AA1-AA17) is given in Table 1 and Table 2.

The CAZy database contained entries with identical sequence IDs and organisms for the JGI data, leading to redundancy. Duplicate entries were removed. Sequences containing ambiguous residues (’X’) were also eliminated, and those labeled as partial in the header were excluded to maintain dataset integrity.

After preprocessing, each AA1–AA17 family contained distinct sets of sequences. The removal of low quality sequences from positive as well as negative sets significantly reduced the total number of sequences in each family. To remove redundancy and ensure more balanced training and validation sets, we applied CD-HIT with a 70% sequence identity threshold to all families. This step further reduced sequence counts across the dataset. After CD-HIT filtering, the number of sequences that remained for AA1-AA17 ranged between 11.5% to 51%.

A machine learning-based model was developed for all eight AA families of LPMOs as classified by CAZy (AA9-AA11 and AA13-AA17). The amount of sequence data available and the resulting sequences from the cleaned and CD-HIT-filtered datasets were carefully checked and curated to ensure suitability for machine learning training. To optimize model training, we divided the data into training and test sets using a 60:40 ratio. To prevent the model’s predictive performance from being skewed by potential biases induced by imbalanced data, we maintained a near 1:1 ratio between positive and negative samples during training set construction (Table 5).

### Feature identification and selection

Feature sets were generated using iFeature in Python [13]. iFeature produced 21 descriptor sets totaling 13494 features per sequence. The R package, Peptides (https://cran.r-project.org/web/packages/Peptides/index.html), contributed an additional 6 features, resulting in 13500 features (Supplementary Table 2).

We selected significant features by implementing two separate ensemble feature selection approaches using Python code for (1) binary and for (2) multiclass models, separately. In both approaches, values were assigned to each descriptor, ranging from 0.1 to 1 where a value closer to 1 is highly significant for a descriptor. We set two thresholds (0.5 and 0.6) to identify the minimal set of features to train a model. Features with values of 0.5 and above were selected. At the 0.5 threshold, none of the features generated by Peptides met the criteria. So, we selected features derived from the iFeature tool that were above the 0.5 threshold value. The selected feature sets included: composition of k-spaced amino acid pairs (CKSAAP, 2,400 descriptors), tripeptide composition (TPC, 8,000 descriptors), amino acid composition (AAC, 20 descriptors), distribution (CTDD, 195 descriptors) and pseudo-amino acid composition (PAAC, 50 descriptors). These features were selected across the families.

### Binary model performance

The features selected were used as input into machine learning algorithms, such as SGD classifiers with log loss, hinge, and modified Huber, as well as neural network (NN) models.

Hyperparameter optimization for SGD and NN was conducted using GridSearchCV in Python. Throughout the training, candidate models were fitted using different parameter combinations, and their performance was evaluated using scoring metrics on validation data to identify optimal configurations.

As noted in previous studies, feature generation methods can sometimes lead to a loss of critical information.^27,28^ To overcome this limitation, we implemented Bi-LSTM models, which can directly learn abstract sequence patterns from raw protein sequences and are well-suited for capturing long-range dependencies. The capability of LSTMs to retain information from more than 1,000-time steps has led to their wide application.^29,30^

All binary classification models were trained and evaluated using cleaned datasets as well as datasets filtered with CD-HIT at a 70% sequence identity threshold to evaluate the effect of redundancy reduction. Across all LPMO families, the Bi-LSTM model consistently achieved higher F1-scores than the feature-based SGD and NN models.

### Multiclass model performance

In addition to developing binary classifiers, we also trained a single multiclass model capable of distinguishing all eight LPMO families. For this multiclass approach, we compared feature-based machine learning methods, such as SGD as well as deep learning architectures like recurrent neural networks (RNN) and Bi-LSTM.

These models were trained and tested using the cleaned as well as the CD-HIT non redundant datasets. This multiple approach enabled us to systematically compare the performance of feature-based and deep learning approaches across binary and multiclass classification tasks to assess the impact of redundancy reduction on the robustness of classifiers for LPMO family prediction. The evaluation metrics of the multiclass models is listed in Table 6.

**Table 6:** Comparison of the evaluation metrics of the multiclass approach of the models, SGD, feature-based NN, Bi-LSTM and dbCAN3 on validation as well as independent set.

| Family | Approach |  | Stochastic gradient descent |  |  | Feature-based neural network | Recurrent Neural Network | dbCAN<sub>3</sub> |
| --- | --- | --- | --- | --- | --- | --- | --- | --- |
|  | Variant |  | Hinge | Log | Modified Huber |  | Bi-Long Short-Term Memory | HMMer, DIAMOND |
| AA<sub>9</sub> | Validation Set | True Positive | 5312 | 5158 | 5229 | 4600 | 5854 | 5846 |
|  |  | False Positive | 1095 | 626 | 597 | 607 | 251 | 0 |
|  |  | False Negative | 676 | 830 | 759 | 1388 | 134 | 49 |
|  |  | True Negative | 19869 | 20338 | 20362 | 20457 | 20711 | 0 |
|  |  | Recall | 0.89 | 0.86 | 0.87 | 0.9372 | 0.977622 | 0.9917 |
|  |  | Precision | 0.83 | 0.91 | 0.9 | 0.7682 | 0.958886 | 1 |
|  |  | F1-score | 0.86 | 0.89 | 0.89 | 0.8443 | 0.968163 | 0.9958 |
|  | Independent Set | True Positive | 104 | 100 | 101 | 91 | 114 | 112 |
|  |  | False Positive | 114 | 19 | 29 | 6 | 7 | 0 |
|  |  | False Negative | 10 | 14 | 13 | 23 | 0 | 1 |
|  |  | True Negative | 1356 | 1451 | 1441 | 1464 | 1462 | 0 |
|  |  | Recall | 0.91 | 0.88 | 0.89 | 0.9381 | 0.9421 | 0.9912 |
|  |  | Precision | 0.48 | 0.84 | 0.78 | 0.7982 | 1 | 1 |
|  |  | F1-score | 0.63 | 0.86 | 0.83 | 0.8626 | 0.9702 | 0.9956 |
| AA<sub>10</sub> | Validation Set | True Positive | 2480 | 2735 | 2500 | 4131 | 4434 | 4592 |
|  |  | False Positive | 1314 | 1837 | 1665 | 2748 | 50 | 0 |
|  |  | False Negative | 2152 | 1897 | 2132 | 501 | 198 | 25 |
|  |  | True Negative | 21006 | 20383 | 20650 | 19572 | 22268 | 0 |
|  |  | Recall | 0.54 | 0.59 | 0.54 | 0.6005 | 0.957254 | 0.9946 |
|  |  | Precision | 0.65 | 0.6 | 0.6 | 0.8918 | 0.988849 | 1 |
|  |  | F1-score | 0.59 | 0.59 | 0.57 | 0.7177 | 0.972795 | 0.9973 |
|  | Independent Set | True Positive | 186 | 213 | 189 | 358 | 422 | 414 |
|  |  | False Positive | 185 | 157 | 150 | 268 | 3 | 0 |
|  |  | False Negative | 239 | 212 | 236 | 67 | 2 | 4 |
|  |  | True Negative | 974 | 1002 | 1009 | 891 | 1156 | 0 |
|  |  | Recall | 0.44 | 0.5 | 0.44 | 0.5719 | 0.9929 | 0.9904 |
|  |  | Precision | 0.66 | 0.58 | 0.56 | 0.8424 | 0.9953 | 1 |
|  |  | F1-score | 0.53 | 0.54 | 0.49 | 0.6813 | 0.9941 | 0.9952 |
| AA11 | Validation Set | True Positive | 1257 | 1264 | 1252 | 1275 | 1360 | 1357 |
|  |  | False Positive | 244 | 219 | 214 | 94 | 18 | 0 |
|  |  | False Negative | 120 | 113 | 125 | 102 | 17 | 5 |
|  |  | True Negative | 25331 | 25356 | 25356 | 25481 | 25555 | 0 |
|  |  | Recall | 0.91 | 0.92 | 0.91 | 0.9313 | 0.987654 | 0.9963 |
|  |  | Precision | 0.84 | 0.86 | 0.85 | 0.9259 | 0.986938 | 1 |
|  |  | F1-score | 0.87 | 0.89 | 0.88 | 0.9286 | 0.987296 | 0.9982 |
|  | Independent Set | True Positive | 41 | 41 | 43 | 43 | 41 | 43 |
|  |  | False Positive | 15 | 17 | 20 | 3 | 1 | 0 |
|  |  | False Negative | 2 | 2 | 0 | 0 | 2 | 0 |
|  |  | True Negative | 1526 | 1524 | 1521 | 1538 | 1539 | 0 |
|  |  | Recall | 0.95 | 0.95 | 1 | 0.9348 | 0.9762 | 1 |
|  |  | Precision | 0.73 | 0.71 | 0.68 | 1 | 0.9535 | 1 |
|  |  | F1-score | 0.83 | 0.81 | 0.81 | 0.9663 | 0.9647 | 1 |
| AA13 | Validation Set | True Positive | 145 | 144 | 145 | 148 | 154 | 156 |
|  |  | False Positive | 141 | 17 | 148 | 70 | 2 | 0 |
|  |  | False Negative | 16 | 17 | 16 | 13 | 7 | 0 |
|  |  | True Negative | 26650 | 26774 | 26638 | 26721 | 26787 | 0 |
|  |  | Recall | 0.9 | 0.89 | 0.9 | 0.6789 | 0.956522 | 1 |
|  |  | Precision | 0.51 | 0.51 | 0.49 | 0.9193 | 0.987179 | 1 |
|  |  | F1-score | 0.65 | 0.65 | 0.64 | 0.781 | 0.971609 | 1 |
|  | Independent Set | True Positive | 1 | 1 | 1 | 2 | 2 | 2 |
|  |  | False Positive | 2 | 2 | 3 | 0 | 0 | 0 |
|  |  | False Negative | 1 | 1 | 1 | 0 | 0 | 0 |
|  |  | True Negative | 1580 | 1580 | 1579 | 1582 | 1582 | 0 |
|  |  | Recall | 0.5 | 0.5 | 0.5 | 1 | 1 | 1 |
|  |  | Precision | 0.33 | 0.33 | 0.25 | 1 | 1 | 1 |
|  |  | F1-score | 0.4 | 0.4 | 0.33 | 1 | 1 | 1 |
| AA14 | Validation Set | True Positive | 497 | 498 | 493 | 514 | 543 | 548 |
|  |  | False Positive | 303 | 185 | 270 | 111 | 6 | 0 |
|  |  | False Negative | 58 | 57 | 62 | 41 | 12 | 2 |
|  |  | True Negative | 26094 | 26212 | 26122 | 26286 | 26389 | 0 |
|  |  | Recall | 0.9 | 0.9 | 0.89 | 0.8224 | 0.978378 | 0.9964 |
|  |  | Precision | 0.62 | 0.67 | 0.65 | 0.9261 | 0.989071 | 1 |
|  |  | F1-score | 0.73 | 0.77 | 0.75 | 0.8712 | 0.983696 | 0.9982 |
|  | Independent Set | True Positive | 19 | 17 | 18 | 21 | 20 | 21 |
|  |  | False Positive | 20 | 28 | 31 | 26 | 2 | 0 |
|  |  | False Negative | 4 | 6 | 5 | 2 | 3 | 0 |
|  |  | True Negative | 1541 | 1533 | 1530 | 1535 | 1558 | 0 |
|  |  | Recall | 0.83 | 0.74 | 0.78 | 0.4468 | 0.9091 | 1 |
|  |  | Precision | 0.32 | 0.38 | 0.37 | 0.913 | 0.8696 | 1 |
|  |  | F1-score | 0.46 | 0.5 | 0.5 | 0.6 | 0.8889 | 1 |
| AA15 | Validation Set | True Positive | 169 | 168 | 173 | 232 | 246 | 261 |
|  |  | False Positive | 342 | 101 | 276 | 355 | 16 | 0 |
|  |  | False Negative | 100 | 101 | 96 | 37 | 22 | 5 |
|  |  | True Negative | 26341 | 26482 | 26402 | 26328 | 26666 | 0 |
|  |  | Recall | 0.63 | 0.62 | 0.64 | 0.3952 | 0.91791 | 0.9812 |
|  |  | Precision | 0.33 | 0.38 | 0.38 | 0.8625 | 0.938931 | 1 |
|  |  | F1-score | 0.43 | 0.47 | 0.48 | 0.5421 | 0.928302 | 0.9905 |
|  | Independent Set | True Positive | 26 | 26 | 26 | 27 | 29 | 29 |
|  |  | False Positive | 19 | 15 | 17 | 17 | 1 | 0 |
|  |  | False Negative | 3 | 3 | 3 | 2 | 0 | 0 |
|  |  | True Negative | 1536 | 1540 | 1538 | 1538 | 1553 | 0 |
|  |  | Recall | 0.9 | 0.9 | 0.9 | 0.551 | 0.9667 | 1 |
|  |  | Precision | 0.59 | 0.63 | 0.6 | 0.931 | 1 | 1 |
|  |  | F1-score | 0.71 | 0.74 | 0.72 | 0.6923 | 0.9831 | 1 |
| AA16 | Validation Set | True Positive | 368 | 367 | 369 | 404 | 422 | 427 |
|  |  | False Positive | 161 | 108 | 212 | 80 | 7 | 0 |
|  |  | False Negative | 68 | 69 | 67 | 32 | 14 | 2 |
|  |  | True Negative | 26355 | 26308 | 26299 | 26436 | 26507 | 0 |
|  |  | Recall | 0.84 | 0.84 | 0.85 | 0.8347 | 0.96789 | 0.9953 |
|  |  | Precision | 0.7 | 0.65 | 0.64 | 0.9266 | 0.983683 | 1 |
|  |  | F1-score | 0.76 | 0.73 | 0.73 | 0.8783 | 0.975723 | 0.9977 |
|  | Independent Set | True Positive | 4 | 4 | 5 | 5 | 5 | 5 |
|  |  | False Positive | 7 | 10 | 9 | 2 | 1 | 0 |
|  |  | False Negative | 1 | 1 | 0 | 0 | 0 | 0 |
|  |  | True Negative | 1572 | 1569 | 1570 | 1577 | 1577 | 0 |
|  |  | Recall | 0.8 | 0.8 | 1 | 0.625 | 0.8333 | 1 |
|  |  | Precision | 0.25 | 0.29 | 0.36 | 1 | 1 | 1 |
|  |  | F1-score | 0.38 | 0.42 | 0.53 | 0.7692 | 0.9091 | 1 |
| AA17 | Validation Set | True Positive | 3 | 4 | 0 | 31 | 51 | 57 |
|  |  | False Positive | 69 | 61 | 156 | 806 | 7 | 0 |
|  |  | False Negative | 55 | 54 | 53 | 27 | 6 | 0 |
|  |  | True Negative | 26825 | 26733 | 26738 | 26088 | 26886 | 0 |
|  |  | Recall | 0.05 | 0.07 | 0.09 | 0.037 | 0.894737 | 1 |
|  |  | Precision | 0.04 | 0.02 | 0.03 | 0.5345 | 0.87931 | 1 |
|  |  | F1-score | 0.05 | 0.04 | 0.05 | 0.0693 | 0.886957 | 1 |
|  | Independent Set | True Positive | 0 | 0 | 0 | 0 | 0 | 0 |
|  |  | False Positive | 0 | 0 | 0 | 0 | 0 | 0 |
|  |  | False Negative | 0 | 0 | 0 | 0 | 0 | 0 |
|  |  | True Negative | 1584 | 1584 | 1584 | 1584 | 1584 | 0 |
|  |  | Recall | 0 | 0 | 0 | 0 | 0 | 0 |
|  |  | Precision | 0 | 0 | 0 | 0 | 0 | 0 |
|  |  | F1-score | 0 | 0 | 0 | 0 | 0 | 0 |

Among the feature-based models, both SGD and feature-based NN classifiers showed variable performance across LPMO families. These models performed reasonably well for well-represented families such as AA9 and AA10 but showed reduced accuracy for smaller families, suggesting that feature-based representations alone may not fully capture the sequence patterns needed for reliable family-level classification.

The Bi-LSTM model achieved higher accuracy and F1-scores for LPMO family classification, with validation F1-scores ranging from 0.9905 to 0.9982 and independent set F1-scores from 0.9952 to 1.0 for most families. The performance of the model is illustrated by the confusion matrix (Supplementary Figure 1), which highlights the model’s consistent and robust predictions across all classes.

### Model evaluation and validation

To evaluate the performance of the model, we used an independent dataset of sequences from CAZy. This dataset included sequences with available annotations spanning multiple LPMO families, with the number of sequences per family ranging from 2 to 425 (Table 6), reflecting substantial class imbalance.

The Bi-LSTM model demonstrated the best predictive performance on the independent dataset, achieving an overall accuracy of 99% and a weighted average F1-score of 0.99 (Table 6). The major LPMO families, AA9 and AA10, showed near-perfect precision and recall (0.94-1.00 and 0.99, respectively), indicating reliable classification for well-represented groups. The model also performed robustly for smaller classes, such as AA14 (F1-score 0.89) and AA16 (F1-score 0.91). The performance was somewhat lower for these rare families due to limited sample size. Notably, non-LPMO sequences (AA1-AA8, AA12) were identified with perfect precision and 99% recall, minimizing false positives.

Multiclass precision-recall curves (Figure 4A) illustrate the relationship between precision and recall for each class across varying thresholds, providing insight into the classifier’s ability to correctly identify positive instances in the presence of class imbalance. Most LPMO families maintained high precision across a broad range of recall values, indicating robust positive predictive value and sensitivity. Multiclass receiver operating characteristic (ROC) curves (Figure 4B) further demonstrate the model’s discriminative capability, with all classes achieving areas under the curve close to or equal to 1.00, signifying excellent separability between classes. Together, the precision-recall and ROC analyses demonstrate that the Bi-LSTM model maintains high sensitivity and specificity across thresholds, with minimal misclassification.

**Figure 4.**
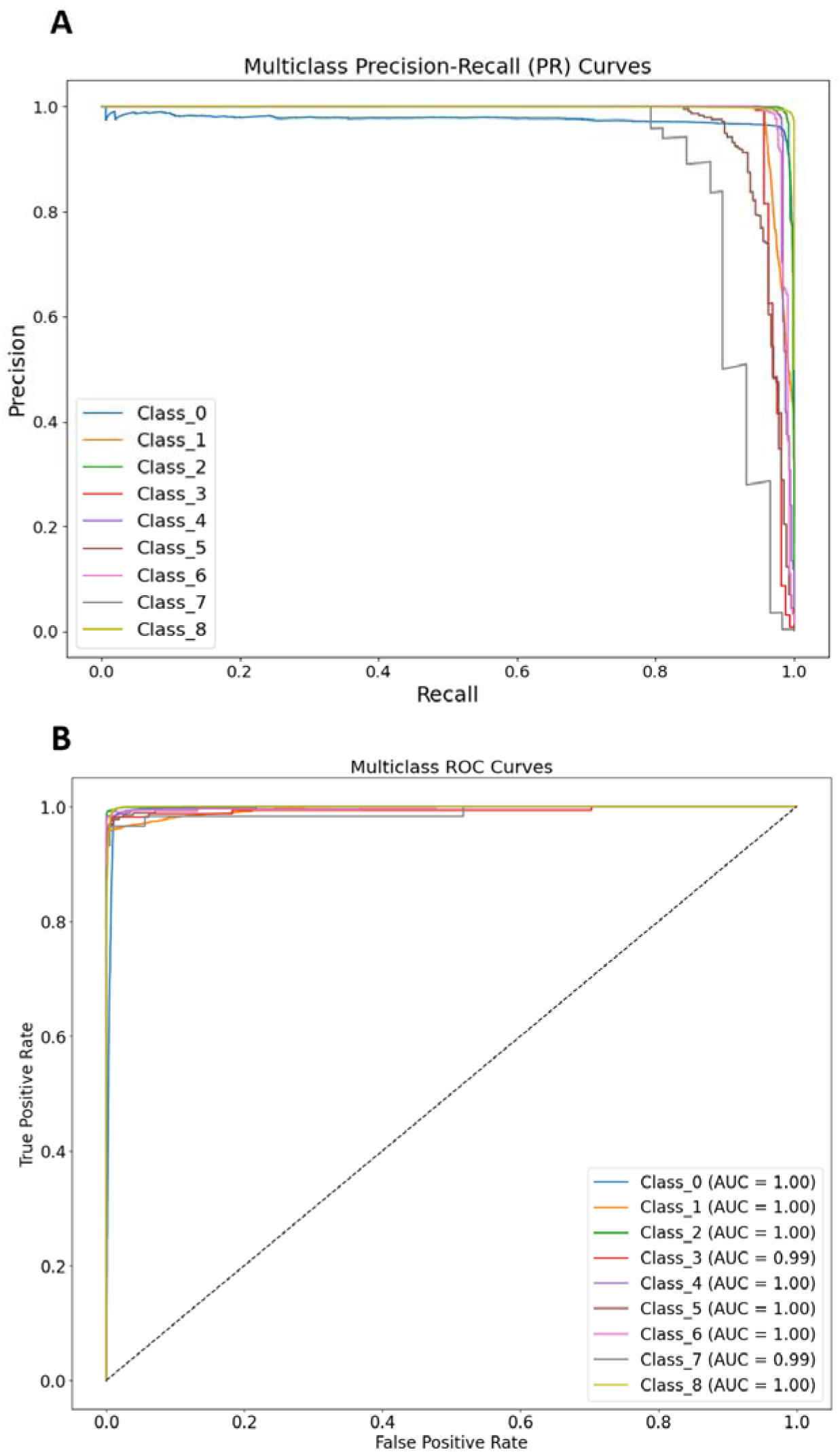
Precision-Recall (PR) and Receiver Operating Characteristic (ROC) Curves of the Bi-LSTM Model. (A) PR curves illustrate the relationship between precision and recall for each class. (B) ROC curves depict the relationship between true positive rate and false positive rate. In both panels, each colored line represents a different class (Class 0: AA9, Class 1: AA10, Class 2: AA11, Class 3: AA13, Class 4: AA14, Class 5: AA15, Class 6: AA16, Class 7: AA17, Class 8: a composite of AA1-AA8, and AA12) collectively demonstrating the model’s discriminative performance across all classes.

In comparison, dbCAN3 achieved consistently high performance across all evaluated LPMO families, with F1-scores ranging from 0.9958 to 1.0000. Among machine learning–based approaches, the Bi-LSTM model achieved performance comparable to dbCAN3, whereas the feature-based models exhibited greater variability across families (F1-scores ranging from 0.0693 to 1.0000), with only the AA14 family approaching dbCAN3-level accuracy.

### PreDSLpmo v2 identifies novel LPMO-like sequences beyond classical UniProt, Pfam and dbCAN3 annotations

To evaluate PreDSLpmo v2.0 against traditional protein annotation methods for identifying LPMOs, we compared each family in the independent dataset against Pfam,^31^ dbCAN3, and BlastP, using the UniProtKB/swissprot database as the search target. Out of 641, Pfam identified 552, dbCAN3 identified 632 and BlastP identified 419 potential LPMOs whereas our deep learning model demonstrated higher sensitivity. PreDSLpmo V2.0 detected 635 candidate LPMOs, including some previously uncharacterized hypothetical proteins that had escaped annotation in existing reference databases. The confusion matrix built for this analysis is given in Table 7.

**Table 7:** Confusion-matrix summary for all AA families merged as one LPMO class. TP: correct LPMO hits for each LPMO family; FP: non-LPMO mislabeled as LPMO; FN: missed LPMO for each family; TN: correct non-LPMO calls. The table also shows precision, recall, and F1-score derived from these counts.

| Tool | TP | FP | FN | TN | Precision | Recall | F1- Score |
| --- | --- | --- | --- | --- | --- | --- | --- |
| BlastP | 419 | 0 | 222 | 0 | 1 | 0.654 | 0.791 |
| Pfam | 552 | 0 | 89 | 0 | 1 | 0.861 | 0.925 |
| dbCAN3 | 632 | 0 | 9 | 0 | 1 | 0.986 | 0.993 |
| PreDSLPMO | 635 | 0 | 6 | 0 | 1 | 0.991 | 0.995 |

### Comparative analysis of PreDSLpmo V2.0 and dbCAN3 predictions

The model was trained on data available before 26 January 2025. To verify whether the model could perform reliably on a newly identified dataset, we collected the sequences of 3,086 proteins, deposited between 1 February and 30 April 2025. After removing 83 redundant entries, we used dbCAN3 to analyze 3,003 unique proteins. Of these, 84 sequences were not predicted. Apart from these 84 unassigned proteins, there were 11 additional sequences that clearly belonged to other CAZyme families (CBM73, GH43_18, GH63, GH18, CBM20, GH1). These were excluded from further LPMO analysis due to their unrelated functions. The remaining 2,908 sequences were annotated as LPMOs by both dbCAN3 and PreDSLpmo.

Within the 84 dbCAN3-unassigned proteins, PreDSLpmo classified 53 as putative LPMOs and 31 as non-LPMO sequences. All 53 candidates, predicted as LPMOs, were examined using Pfam and CDD to check for recognizable LPMO-related domains. Most sequences showed domain signatures consistent with known LPMO families, including AA9 and AA10/LPMO10 motifs, BIM1-like or LPMO_auxiliary-like copper-associated regions, DOMON/DOH_DOMON monooxygenase domains, and CBD9-like carbohydrate-binding modules. Only a single sequence lacked interpretable domains. Although these domain signals were too weak or fragmented for dbCAN3 to classify, their presence supports the biological relevance of PreDSLpmo predictions. This result suggests that PreDSLpmo can complement existing annotation pipelines by expanding the identification of candidate LPMOs beyond those captured by dbCAN3.

All 53 candidates predicted by PreDSLpmo were analyzed using SignalP,^32^ which predicts the presence of Sec/SPI signal peptides and their cleavage positions. SignalP identified secretion signals in 34 sequences. In these cases, the predicted cleavage site was immediately followed by an N-terminal histidine, a hallmark of the LPMO histidine brace formed by the N-terminal His and the internal His residue. This arrangement, commonly reported in experimentally characterized LPMOs, supports the likelihood that these candidates represent functional or functionally related LPMO homologs.

The evidence from the Pfam/CDD domains and from the SignalP secretion profiles provides independent support for the reliability of the LPMO classifications generated by PreDSLpmo. The presence of LPMO-associated domains indicates that most PreDSLpmo identified candidates are biologically credible LPMO family members. These findings show that PreDSLpmo can recover divergent or weakly conserved LPMOs that classical HMM-based pipelines fail to annotate, demonstrating the added biological and functional resolution provided by the deep-learning framework. Nevertheless, these computational predictions should be complemented with experimental validation to ensure greater accuracy.

### PreDSLpmo web server

We implemented PreDSLpmo V2.0, the multiclass trained model based on Bi-LSTM, as a web server. The web server, PredLpmo, addresses the critical need for accessible, high-throughput prediction platforms capable of accurately distinguishing between LPMO families and non-LPMO sequences as depicted in Figure 5. The homepage (Figure 5A) introduces users to the server’s deep learning-based prediction capabilities, emphasizing its automated workflow and support for high-throughput LPMO classification and sequence analysis across diverse auxiliary activity (AA) families.

**Figure 5:**
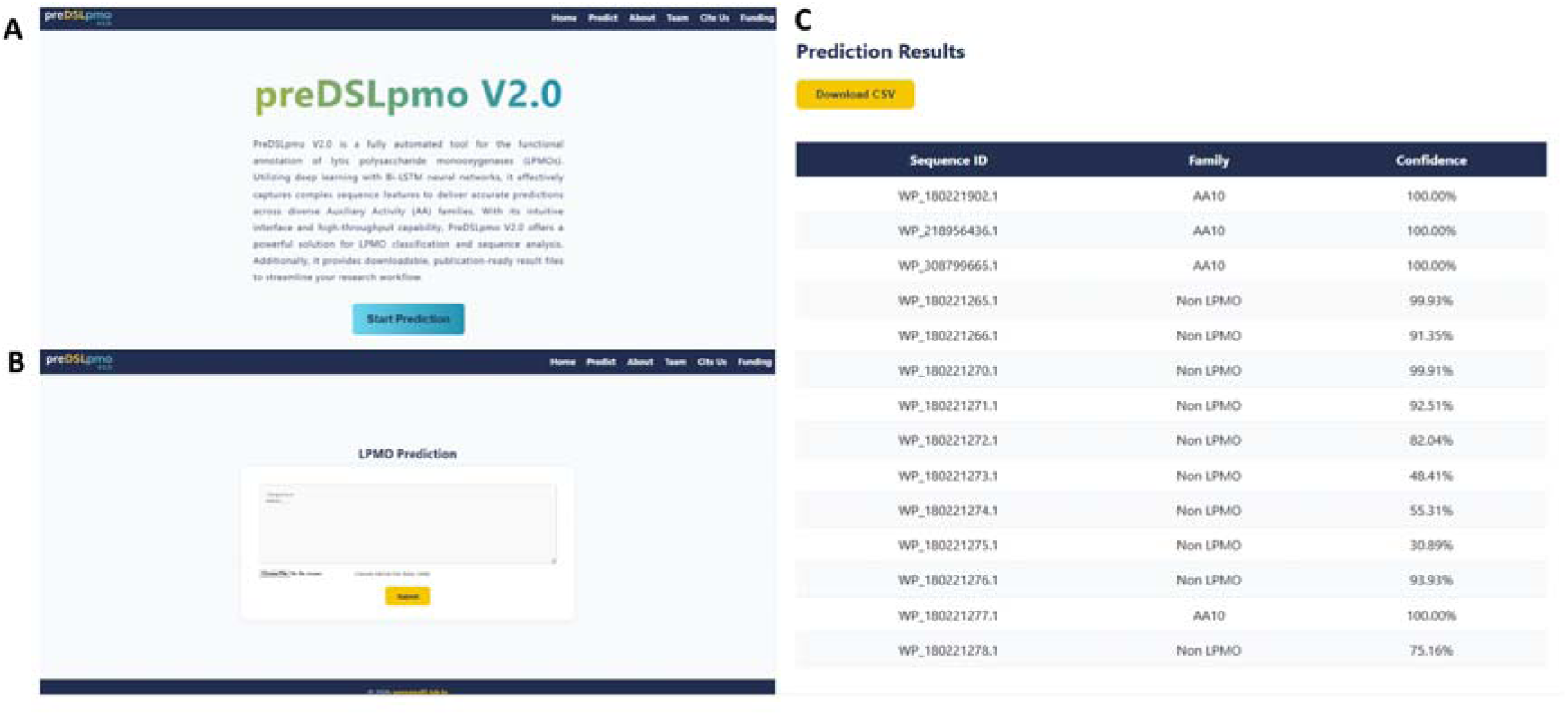
Overview of the PreDSLpmo V2.0 web server interface and workflow. (A) Homepage of the PreDSLpmo V2.0 web server, highlighting its deep learning-based prediction capabilities, user-friendly design, and options for high-throughput LPMO classification and sequence analysis. (B) LPMO prediction submission page, where users can input protein sequences in FASTA format directly into the text box or upload a file for automated LPMO classification and analysis. (C) Prediction results for *Streptomyces* sp. UH6 protein sequences showing classification as LPMOs or Non-LPMO proteins. Each entry lists the sequence ID, predicted family, and model confidence score. Results can be exported as a CSV file for further analysis.

The submission page (Figure 5B) allows users to input protein sequences either by pasting them in FASTA format or by uploading a file, facilitating flexible and efficient data entry for automated analysis. Upon submission, the results page (Figure 5C) presents the prediction outcomes in a clear tabular format, displaying the sequence ID, predicted family classification as either belonging to a specific AA family or as belonging to the non-LPMO class. For each prediction, the results provide an associated confidence score for each input sequence. Additionally, users can download the results as a CSV file for further analysis, for individual as well as high-throughput protein annotation tasks.

## Conclusion

This study demonstrates how sequence-based deep learning can be used for the reliable family level classification of LPMOs. Comparing multiple machine learning methods, we found that the multiclass Bi-LSTM model consistently provided accurate and stable predictions across validation and independent datasets. The Bi-LSTM model highlights the value of directly learning from raw protein sequences, enabling the capture of long-range dependencies and family-specific sequence patterns that are difficult to represent using only handcrafted features. The model maintained high accuracy for well-represented LPMO families and showed robust performance even for smaller families, despite limited sample sizes. These results support the reliability of sequence-based deep learning for enzyme family classification under realistic and imbalanced data conditions.

The robust classification of LPMO families has important practical implications for downstream research. Reliable computational predictions can assist in prioritizing candidate enzymes for biochemical characterization, guide the formulation of enzyme cocktails for biomass degradation, and support the discovery of LPMOs from rapidly expanding sequence databases. By reducing false positives and providing clear family assignments, the multiclass approach supports large-scale sequence screening.

The optimized Bi-LSTM model has been deployed as the PreDSLpmo v2.0 web server (https://predlpmo.in), enabling researchers to submit raw protein sequences and obtain predicted LPMO family assignments along with confidence scores. Our model can be independently tested by researchers worldwide for classifying AA family sequences added after May 2025.

The Bi-LSTM model and the training, testing, and validation pipeline adopted for AA family classification can also be transposed for use in the classification of other protein families. Collectively, this work bridges the gap between large-scale sequence data and experimental enzymology, providing a practical computational resource for bioenergy research and carbohydrate-active enzyme discovery.

## CRediT authorship contribution statement

Sweety Deena Ramesh: Writing – original draft, writing, review and editing, methodology, data curation, validation, website development. Kavin S Arulselvan: Writing – original draft, methodology, data curation, visualization, website development. Vaishnavi Saravanan: Writing, review and editing, data curation, visualization, website development. Priyadharshini Pulavendran: Writing, review and editing, data curation, visualization, website development. Ragothaman M. Yennamalli: Supervision, conceptualization, funding acquisition.

## Acknowledgements

All the authors thank SASTRA Deemed to be University for infrastructural support. RMY was funded by the Department of Biotechnology, Ministry of Science and Technology, Government of India (BT/PR42225/BCE/8/1586/2021)

## Code Availability

The codes used in this study are available at: https://github.com/raghuyennamalli/PreDSLpmo_v2 and the webserver is available at: https://predlpmo.in/

