## Supplementary Tables for "Sequence-based deep learning model for annotating lytic polysaccharide monooxygenase families"

**Supplementary Table 1:** EC Numbers of AA Families (LPMO and Non-LPMO).

| **Auxiliary Activity**  **LPMO Family** | **EC Number** | **Auxiliary Activity NON-LPMO**  **Family** | **EC Number** |
| --- | --- | --- | --- |
| AA9 | EC 1.14.99.54  EC 1.14.99.56  EC 1.14.99  EC 1.14.99 | AA1 | EC [1.10.3](http://www.enzyme-database.org/query.php?ec=1.10.3.-)  EC [1.10.3.2](http://www.enzyme-database.org/query.php?ec=1.10.3.2) |
| AA10 | EC 1.14.99.53  EC 1.14.99.54  EC 1.14.99.56  EC 1.14.99 | AA2 | EC [1.11.1](http://www.enzyme-database.org/query.php?ec=1.11.1.-)  EC [1.11.1.11](http://www.enzyme-database.org/query.php?ec=1.11.1.11)  EC [1.11.1.13](http://www.enzyme-database.org/query.php?ec=1.11.1.13)  EC [1.11.1.14](http://www.enzyme-database.org/query.php?ec=1.11.1.14)  EC [1.11.1.16](http://www.enzyme-database.org/query.php?ec=1.11.1.16)  EC [1.11.1.5](http://www.enzyme-database.org/query.php?ec=1.11.1.5) |
| AA11 | EC 1.14.99.53 | AA3 | EC [1.1.3.10](http://www.enzyme-database.org/query.php?ec=1.1.3.10)  EC [1.1.3.13](http://www.enzyme-database.org/query.php?ec=1.1.3.13)  EC [1.1.3.16](http://www.enzyme-database.org/query.php?ec=1.1.3.16)  EC [1.1.3.4](http://www.enzyme-database.org/query.php?ec=1.1.3.4)  EC [1.1.3.7](http://www.enzyme-database.org/query.php?ec=1.1.3.7)  EC [1.1.5](http://www.enzyme-database.org/query.php?ec=1.1.5.-)  EC [1.1.5.9](http://www.enzyme-database.org/query.php?ec=1.1.5.9)  EC [1.1.99.18](http://www.enzyme-database.org/query.php?ec=1.1.99.18)  EC [1.1.99.29](http://www.enzyme-database.org/query.php?ec=1.1.99.29) |
| AA13 | EC 1.14.99.55 | AA4 | EC [1.1.3.38](http://www.enzyme-database.org/query.php?ec=1.1.3.38) |
| AA14 | EC 1.14.99 | AA5 | EC [1.1.3](http://www.enzyme-database.org/query.php?ec=1.1.3.-)  EC [1.1.3.13](http://www.enzyme-database.org/query.php?ec=1.1.3.13)  EC [1.1.3.47](http://www.enzyme-database.org/query.php?ec=1.1.3.47)  EC [1.1.3.7](http://www.enzyme-database.org/query.php?ec=1.1.3.7)  EC [1.1.3.9](http://www.enzyme-database.org/query.php?ec=1.1.3.9),  EC [1.2.3.15](http://www.enzyme-database.org/query.php?ec=1.2.3.15) |
| AA15 | EC 1.14.99.53  EC 1.14.99.54 | AA6 | EC [1.6.5.6](http://www.enzyme-database.org/query.php?ec=1.6.5.6) |
| AA16 | EC 1.14.99.54 | AA7 | EC 1.1.3  EC 1.1.99 |
| AA17 | EC 1.14.99 | AA8 | No protein characterized |
|  | | AA12 | EC 1.-.-.- |

**Supplementary Table 2**: The total number of features extracted from python iFeature package.

| **Packages** | **Descriptors** | **Dimension** |
| --- | --- | --- |
| Python package (iFeature) | Amino Acid Composition (AAC) | 20 |
|  | Composition of k-spaced Amino Acid Pairs (CKSAAP) | 2400 |
|  | Dipeptide Composition (DPC) | 400 |
|  | Dipeptide Deviation from Expected Mean (DDE) | 400 |
|  | Tripeptide Composition (TPC) | 8000 |
|  | Grouped Amino Acid Composition (GAAC) | 5 |
|  | Composition of k-spaced Amino Acid Group Pairs (CKSAAGP) | 150 |
|  | Grouped Dipeptide Composition (GDPC) | 25 |
|  | Grouped Tripeptide Composition (GTPC) | 125 |
|  | Normalized Moreau-Broto (NMBroto) | 240 |
|  | Moran (Moran) | 240 |
|  | Geary (Geary) | 240 |
|  | Composition (CTDC) | 39 |
|  | Transition (CTDT) | 39 |
|  | Distribution (CTDD) | 195 |
|  | Conjoint Triad (CTriad) | 343 |
|  | Conjoint k-spaced Triad (KSCTriad) | 343 |
|  | Sequence-Order-Coupling Number (SOCNumber) | 60 |
|  | Quasi-Sequence-Order Descriptors (QSOrder) | 100 |
|  | Pseudo-Amino Acid Composition (PAAC) | 50 |
|  | Amphiphilic Pseudo-Amino Acid Composition (APAAC) | 80 |
| ‍‍R package (Peptides)  ‍ | Molecular weight | 1 |
|  | Isoelectic point | 1 |
|  | Hydrophobicity | 1 |
|  | Aliphatic index | 1 |
|  | Instability index | 1 |
|  | Boman index | 1 |
| ‍ | **Total number of Features** | **13500** |
