## Supplementary figures and images for "Sequence-based deep learning model for annotating lytic polysaccharide monooxygenase families"

### Supplementary Figures

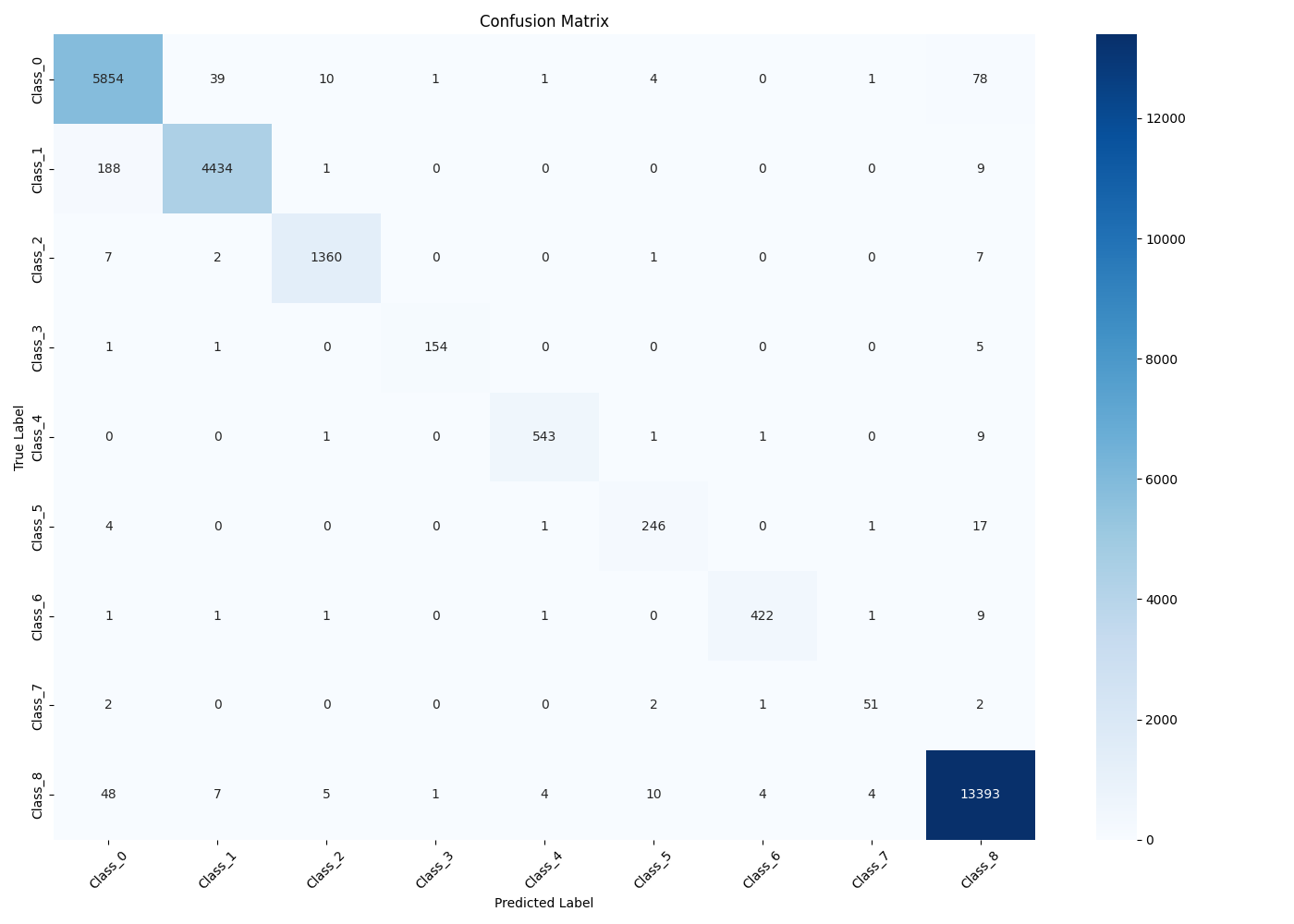

**Supplementary figure 1: Confusion matrix of Bi-LSTM model**
